# Orphan Drugs and Rare Diseases: Unveiling Biological Patterns through Drug Repurposing

**DOI:** 10.1101/2023.05.03.539318

**Authors:** Belén Otero-Carrasco, Santiago Romero-Brufau, Andrea Álvarez-Pérez, Adrián Ayuso-Muñoz, Lucía Prieto-Santamaría, Juan Pedro Caraça-Valente Hernández, Alejandro Rodríguez-González

## Abstract

Rare diseases are a collection of unusual pathologies that afflict millions of individuals globally. However, the creation of treatments for these conditions is frequently limited due to the high expenses and lack of profitability associated with drug development. Orphan drugs, which are medications specifically designed for rare diseases, have played a pivotal role in treating these diseases over the past several years. Nevertheless, their creation remains challenging, and many rare diseases lack approved therapies. Therefore, drug repurposing has emerged as a viable strategy for identifying potential new treatments for these pathologies, a technique consists in using existing drugs to treat a new disease different from the one that they were developed for. This approach can significantly reduce the time and cost of drug development while increasing the likelihood of success. In this paper, we examined the temporal progression of orphan drugs since their introduction and assess the impact of drug repositioning on treatments for rare diseases. Additionally, we aim to identify biological patterns that may be unique to rare diseases treated with repurposed orphan drugs. To this end, we analyzed various biological components associated with these diseases, categorized linked diseases, and obtained the type of orphan drug associated with them. Lastly, we evaluated the phenotypic similarity between diseases treated with an orphan drug through repurposing. Through these findings, we have gained insight into the evolution of orphan drug development in recent years and identified specific patterns that characterize rare diseases associated with them.

## I. Introduction

Rare diseases, also known as orphan diseases, are medical conditions that affect a small percentage of the general population compared to other diseases [1]. The definition of a rare disease varies by country. In the United States, a rare disease is defined as one that affects fewer than 200,000 people. In Europe, a disease is considered rare if it affects fewer than 1 in 2,000 people [2]. According to the World Health Organization (WHO), there are between 6,000 and 8,000 rare diseases, and they affect approximately 300 million people worldwide [3]. In these diseases, genetic factors are estimated to be responsible for 80% of cases [4]. In some parts of the UK, it is estimated that 1 in 17 people will be affected by a rare disease at some point in their lives [5]. The prevalence of rare diseases varies by disease and by region. Some rare diseases are more prevalent in certain ethnic groups or geographic regions [6].

Diagnosing a rare disease is a challenging process that can take anywhere from several months to several decades, contingent upon the patient’s age, phenotype, and access to resources. On average, it takes around 4-5 years to arrive at an accurate diagnosis for a rare disease [7], during which time patients are subjected to a diagnostic journey involving extensive and costly testing. Despite these efforts, patients frequently go undiagnosed or are misdiagnosed [8].

Due to the scarcity of clinical information and inadequate epidemiological data, their prompt diagnosis and effective treatment pose a significant challenge. In an attempt to address this problem, the Orphan Drug Act (ODA) was enacted in the United States in 1983 to address the lack of available treatments for rare diseases [9]. The ODA provides incentives to pharmaceutical companies to develop drugs for rare diseases by offering tax credits, grants, and exclusive marketing rights [10]. This law has had a significant impact on the development of new therapies for rare diseases, with over 600 drugs approved for orphan indications since its enactment [11]. The success of the ODA has led to similar legislation being passed in other countries, such as the Orphan Medicinal Products Regulation in the European Union [12]. Nonetheless, despite the incentives provided by these laws, the number of approved treatments compared to the amount of existing rare diseases is still too limited.

The development of new drugs for rare diseases is encumbered by the significant investment of time and resources required to create a drug de novo, with no guarantee of achieving an effective treatment for the disease under investigation. The introduction of a new compound to the market can incur costs of up to 2.5 billion US dollars, which includes substantial expenses for both development and manufacturing processes [13]. Out of a pool of 5,000 compounds subjected to preclinical testing, on average, only five are advanced to human trials, and ultimately, only one of these five receives therapeutic approval from the US Food and Drug Administration (FDA) [14].

To reduce the time and costs associated with de novo drug development, it is necessary to identify and develop alternatives. In recent years, the pharmaceutical industry has paid increasing attention to drug repurposing (DR) as a cost-effective and rapid method for drug development [15]. Drug repositioning refers to the identification of new therapeutic applications for drugs that have already been approved or are currently in use, but for indications outside their original scope [16]. This approach represents a promising alternative to traditional drug development. The process of repurposing drugs typically takes between 3-12 years for a marketed treatment to reach patients [17]. Repurposed drugs have been found to have a higher success rate of 30% to 75%, with an average cost of $300 million, which is five times less than the cost of developing new drugs [18]. This is especially useful in the context of rare diseases, where DR may offer a faster path to market for potential treatments for the large multitude of rare diseases that lack any associated treatment [2]. Furthermore, the repositioned drug has a pre-existing safety profile and established the pharmacokinetics, which simplifies and expedites the regulatory approval procedure [19].

The evolution of DR is driven by the availability of large-scale omics data and the development of information technology tools to analyze and integrate these data [20]. Drug repositioning represents a promising strategy for addressing the unmet medical needs of patients with rare diseases, particularly in the context of orphan drugs [21]. As the field continues to evolve, it is expected that the identification of new targets and the development of new computational tools will lead to the discovery of even more opportunities for drug repositioning.

## II. Related works

Research on the use of repositioned orphan drugs in rare diseases is scarce. There are extremely few scientific studies that relate the three concepts. Despite this, different investigations that have obtained significant results in this field over the last few years are shown below. One of the most well-known cases of drug repurposing in rare diseases was Thalidomide. This drug was withdrawn from the market due to its teratogenic effects. However, it was later found to be effective for the treatment of multiple myeloma, a rare cancer of plasma cells [22].

Focusing on studies based on computational methods for DR, it is relevant to point out a study that presents a drug repurposing strategy applied to nephropathic cystinosis. This consists of a combination of mechanism-based and cell-based screens that could predict therapeutic responses at both molecular and system levels for identifying potential drugs [23]. Another study indicates the development of a computer program capable of comparing drug binding sites to explore different opportunities for repositioning known drugs to proteins associated with rare diseases [24]. Furthermore, research has been found that uses DR to search for potential new treatments for adrenocortical carcinoma (ACC). They have developed a model called Heter-LP that identifies innovative putative drug-disease, drug-target, and disease-target relationships for this type of cancer [25].

Within DISNET project [26], our knowledge platform for drug repositioning, we have developed a study based on the development of different computational methods to try to obtain potential treatments for a set of 13 rare diseases. Through this study, we were able to obtain a 75% success rate in terms of potential treatments for these diseases [27]. To complement the current line of research, we propose a study that delves into the evolution of drug repositioning for rare diseases over time. Rather than suggesting new potential cases, our study aims to identify patterns that distinguish rare diseases treated with orphan drugs successfully repositioned through the DISNET^1^ platform. This approach will provide us with a better understanding of the relationship between drug repositioning and rare diseases.

## III. Materials and methods

In this section, the methodology used to carry out this study will be explained. A summary of all of the analyses can be seen in Fig 1. First, the data of interest for this study regarding orphan drugs are selected. Then, a temporal and descriptive analysis of the data related to these drugs will be carried out. Finally, we will look for patterns that may be characteristic of the rare diseases related to the orphan drugs studied.

**Fig 1.**
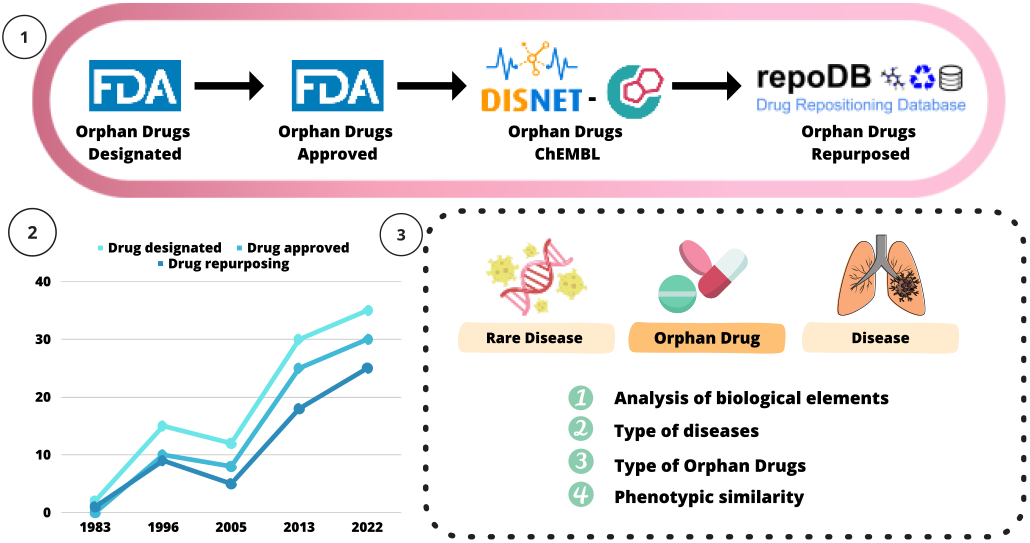
Summary of the different methodologies applied in the present study to understand the orphan drugs evolution and the biological patterns related to rare diseases. The data represented in point 2 are not real data.

### A. Data acquisition and integration

The orphan drugs analyzed in this research were obtained from the FDA’s Orphan Drug Product designation database^2^. On this platform, we found a list of drugs with different associated information. For this study, the attributes that were considered were the following:

- *Generic drug name*: usually refers to the chemical substances that constitute the drug.
- *Trade drug name*: is the name under which the drug has been registered in the market by the pharmaceutical company that developed it.
- *Date designated*: is the date on which the molecule or drug was designated as an orphan drug.
- *Marketing approval date*: is the date on which the drug was approved for marketing.
- *Orphan designation*: corresponds to the description of the rare disease for which an orphan drug has been considered as treatment.
- *Orphan designation status*: the possible status options are Designated, Approved or Withdrawn.

On the FDA website, there are a total of 6,587 drugs that are under consideration to be or have been orphan drugs. These orphan drugs are assigned the category “Designated” in the variable orphan designation status and have a date associated with them in the variable date designated. These data correspond to all the records from the time they began to be considered orphan drugs (01/01/1983) up to the time of the query (03/02/2023). Within them, some cases will be eliminated from the final dataset since, despite being approved by the FDA at the time, they were finally withdrawn as orphan drugs. Removing these cases, and selecting only those with “Approved” designation, the final set of orphan drugs is 1,112 drugs.

Once the orphan drugs approved to treat a certain rare disease were selected, we needed to obtain the CheMBL IDs associated with these drugs to obtain more information about them. To accomplish this task, two different data sources were used. First of them was CheMBL ^3^, which is the primary source of open data that selects and stores bioactivity, molecules, targets, and drugs extracted from multiple repository. The primary relationship captured in the ChEMBL database is the association between a ligand and a biological target [28].

The second database was DISNET. DISNET is a knowledge platform containing different information about biological diseases extracted from public data sources. DISNET is divided into three different layers: (i) the biologic layer that contains information relevant to the diseases such as their genes, (ii) the phenotypic layer that has information related to the symptoms of these diseases, and (iii) the pharmacologic layer that provides information about the treatments of the different diseases. Within this drug layer, among other information, we can find the CheMBL ID associated to each of the registered drugs and previously obtained from CheMBL. By leveraging this platform to identify the CheMBL ID of approved orphan drugs, we extracted a total of 1,051 drug entries that included this information within DISNET.

To conduct this research, it was essential to gather information on the total number of repositioned drugs available up to the present time. In this regard, the list of approved repositioned drugs was obtained through the RepoDB [29] platform. This platform contains a total of 3,757 repositioned drugs. Of these, we obtained only those orphan drugs that had been repositioned and were registered on this database. Of the 1,051 orphan Drugs considered, 365 are repositioned drugs according to the data source consulted.

### B. Temporal exploration of orphan drugs

First analysis carried out in this work focused on the temporal evolution of the orphan drugs, as well as the parallel evolution of the repositioned drugs.

For this purpose, considering the numbers of orphan designated cases and approved cases discussed in the previous section, an average estimate was calculated of how many years it took to approve an orphan drug. To obtain this value, the difference between the date designated and the marketing approval date was calculated for each of the drugs considered. Once these values were obtained for each record, the total average was calculated by time interval. Taking these values into account, the evolution of designated and approved drugs over the years (from the existence of the registers until 2022) was represented temporally. In addition, to this temporal analysis, we added the evolution of drugs repositioned for rare diseases over this time to see how they have been evolving and the importance they are taking in orphan drugs in recent years. With these data, we could get an idea of the future evolution of the relationship between the 3 datasets considered: designated drugs, approved drugs, and repositioned drugs.

### C. Biological patterns associated with rare diseases

Our goal is to identify biological patterns that explain the success of repositioned orphan drugs in treating rare diseases. By analyzing such patterns, we can gain valuable insights into why certain drugs are effective in treating certain rare diseases. This knowledge can serve as a foundation for discovering new repositioning treatments for other rare diseases. Therefore, identifying characteristic patterns associated with rare diseases treated with repositioned drugs can be a promising starting point for future research.

The first step was to understand the rare diseases that are treated through repositioning cases. These rare diseases were registered in the FDA database as broad descriptions. Therefore, it was necessary to obtain an identifying code for each of these diseases. The Unique Concept Identifier (CUI) of the Unified Medical Language System (UMLS)^4^ was used to categorize these descriptions into specific diseases. These CUI codes were obtained through the DISNET platform, which, as mentioned above, contains relevant information on a multitude of diseases, including rare diseases. Associating this code to the orphan designation, a total of 127 repositioning cases of orphan drugs for rare diseases were found. From this total of rare diseases, we obtained the number of genes, symptoms, and pathways associated with each of them, and the resulting mean of the entire dataset was calculated.

Repurposing cases are constructed by two diseases associated with the same drug. This will be referred to as triples throughout the study. In our case, we have the rare disease associated with the orphan drug and one more disease, which can be the original designation of that drug or the new one, since the RepoDB platform does not indicate this relationship. Therefore, we can represent our triple by the following structure: “Rare disease - Orphan drug – Disease”. Bearing this in mind, we focused on finding out what distinguishes the triples that comprise this database. We are interested in knowing whether the two diseases related to the drug are rare or whether only the one associated with the drug in the FDA database is rare. This will allow us to identify whether DR usually happens from one rare disease to another or, on the contrary, from a non-rare disease to a rare disease. The associated disease will be checked for rarity through the Orpha code integrated in the DISNET platform previously extracted from Orphanet^5^. If a relationship is found between the CUI of the disease and an Orphacode, the disease will be considered rare.

Related to the previous point, we want to know what types of diseases are related to the drug. The type of disease will be obtained by using ICD-10-CM (International Classification of Diseases, 10th revision, Clinical Modification) [30] for the rare disease and the disease-associated. In keeping with the current theme, we aim to discern the drug types associated with these rare diseases to determine their most prominent types concerning repurposing for such conditions. To this end, we have considered Anatomical Therapeutic Chemical (ATC) [31] classifications to identify the type of drug. Furthermore, we will combine the data gathered on the disease and drug types in the preceding sections to establish the drug-rare disease pattern, and possibly identify any patterns that link particular disease types with specific drug types.

Finally, we want to calculate the phenotypic similarity between the two diseases that compose the triplet. To have a benchmark, we will compare the similarity that exists between all the diseases contained in the DISNET database. The phenotypic similarity will be calculated using the Jaccard similarity index:

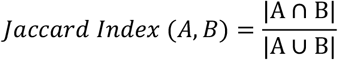

where A and B represent vectors of the biological element studied related to the two diseases that make up the triple, where the presence of that biological element was represented as a 1 and the absence as a 0 in the vector. To test for differences between the similarity values obtained, a Mann-Whitney U-test was applied to the different datasets. This statistical test is used because the data do not follow a normal distribution.

All the procedures carried out to obtain the results shown below by applying the previously mentioned methodologies are available at the following link: https://medal.ctb.upm.es/internal/gitlab/b.otero/orphan_drugs_repurposing

## IV. Results and discussion

### A. Temporal exploration of orphan drugs

In the present study, the aim was to observe the temporal evolution of orphan drugs and repositioned drugs since records have been kept. The orphan drugs, as mentioned in previous sections, first receive the status of designated and after a certain period, only some of them become approved. Therefore, the first thing we have calculated is the mean time it takes to approve a drug on average. This calculation has been made for the total data set as well as for intervals of 13 years since we have records for a total of 39 years. Furthermore, this average approval time was calculated for all approved and repurposed drugs. Table I shows the results of these analyses.

**Table I.** ANALYSIS OF THE ESTIMATED AVERAGE TIME TO APPROVAL OF AN ORPHAN DRUG TAKING INTO ACCOUNT THE DIFFERENT DATASETS STUDIED.

| Years | Mean time to approve orphan drugs |  |
| --- | --- | --- |
|  | Approved drugs | Repurposed drugs |
| 1983 - 1996 | 6.37 | 5.83 |
| 1996 - 2010 | 6.91 | 6.80 |
| 2010 - 2022 | 3.52 | 2.97 |
| Total | 5.39 | 5.35 |

These results show that the average time it takes for an orphan drug to be approved is around 5 years. This average does not vary much between approved and repurposed drugs. It is true that compared to the time it takes for a drug developed de novo to be put on the market, the times are much shorter (∼15 vs ∼5 years). The fact that there is not much difference between approved and repositioned drugs may be largely due to the legal processes required by the different associations that regulate the approval of these drugs.

Once the mean approval time for these drugs had been calculated, we wanted to look more closely at how the designation, approval, and repositioning of orphan drugs have evolved from the time records were kept to the present day. Fig 2 shows the temporal trends of these three datasets. In addition, considering the average approval time, it has been represented how this evolution would be if the designated drug were approved simultaneously and did not have to wait an average of 5 years to reach the market. For this reason, 5 different lines are shown in Fig 2, where Approved/Repurposed_time_ap (time to approved) are plotting it.

**Fig 2.**
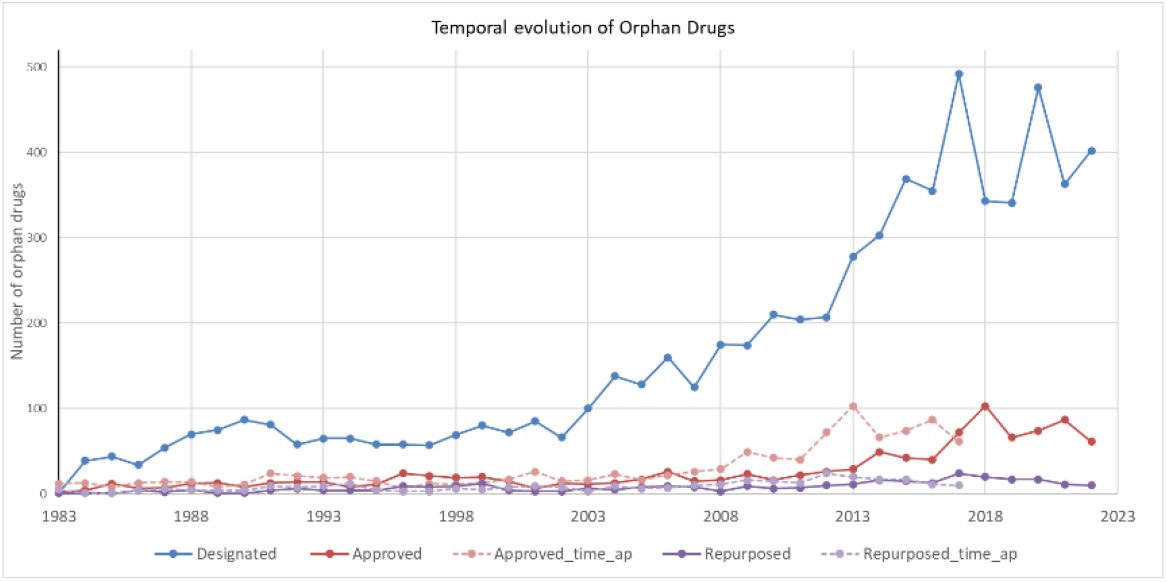
Time representation of the evolution of designated, approved, and repositioned orphan drugs since records have been kept up to the present. In addition, the dashed lines show how this evolution would be if the average approval time were considered.

Through this temporal representation, it has been observed that the involvement of repositioned drugs throughout history seems to have gained more importance. The temporal trend appears to indicate that in the coming years, repositioned drugs will play a more decisive role in being considered as future treatments for various rare diseases. There indeed seems to be a decrease in the number of repositioned cases in the last 2-3 years, but this is accompanied by a decrease in the number of approved drugs. This change in the trend may have been influenced by the years of the COVID-19 pandemic, which may have delayed the approval of several orphan drugs or that more research was done on the use of repositioned drugs for rare diseases.

### B. Biological patterns associated with rare diseases

As part of the search for biological patterns in the rare diseases treated with a repositioned orphan drug, the first result we have obtained concerns the biological elements associated with this group of diseases. In Table II, we can observe the number of genes, symptoms, and pathways (direct and via genes) associated, on average, with the set of rare diseases considered. These results will allow us to know, for future cases of repurposing, the biological characteristics that might be more appropriate for rare diseases to have for successful repositioning to occur.

**Table II.** Descriptive analysis of the biological elements related to rare diseases considering.

| Biological element | Mean | Std <sup>a</sup> |
| --- | --- | --- |
| Genes | 90.63 | 158.64 |
| Pathways direct | 1.20 | 0.44 |
| Pathways via genes | 398.83 | 676.73 |
| Symptoms | 58.62 | 23.12 |

<sup>a</sup>. Standard deviation

Then, it was checked whether the “Rare disease - Orphan drug - Disease” triples were formed by two rare diseases or by one rare and one non-rare disease. Of the total 127 DR cases with orphan drugs, 58 unique triples were obtained. The distribution of these triples according to the criteria was 51 cases of Rare disease – Rare disease (87.9%) and 7 cases of Rare disease – Non-rare disease (12.1%). Therefore, it has been observed that most of the triples are formed by two rare diseases, i.e., both the disease associated with the drug in the FDA database and the other obtained from RepoDB are rare diseases. This is a highly informative fact that indicates that when DR cases in rare diseases occur, in about 90% of the cases, this repurposing is carried out from one rare disease to another. This is a crucial pattern for the search for future DR cases for these pathologies.

In addition, the type of diseases that were composing both sides of the triples and the most common correlation between them was obtained. These results are represented in Fig 3 through a heat map where the dark orange colour indicates a higher relationship between both types of diseases and the lighter colour indicates the opposite case. Observing this figure, the strongest relationship exists between neoplasm-type diseases, where their association value is very close to 1. The appearance of more neoplasm-type triples can be explained because cancers are the most studied type of disease and the development of therapies that can be useful for cancers considered rare is much more abundant than for any other type of disease. Most associations between rare and other diseases occur between diseases of the same type, regardless of whether they appear more or less frequently in the triples considered. Therefore, through these results, we can consider that it is more likely that when a drug is repositioned it is usually between diseases that are categorized under the same class. This notion is highly coherent and valid since diseases of the same category may share more characteristics than in any other case.

**Fig 3.**
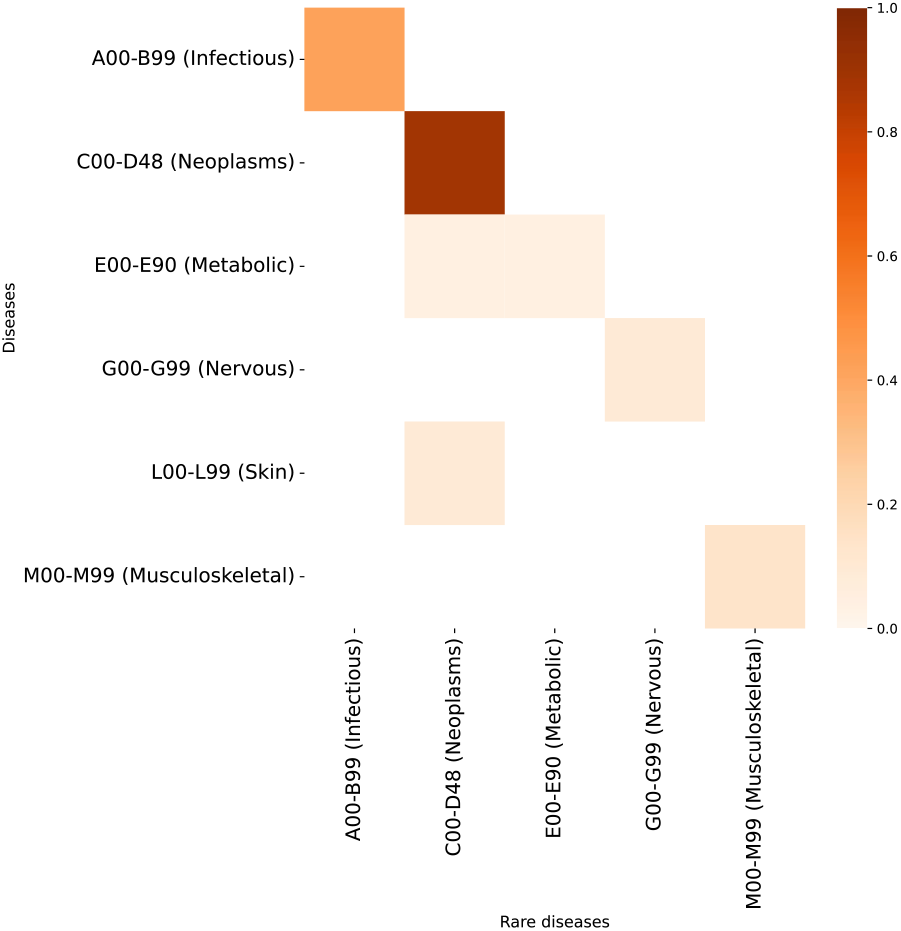
A representation of the relationship between rare diseases and diseases according to the type of disease based on ICD-10-CM.

Once the diseases that make up the triples are known, we have also obtained results on the associated orphan drugs according to the ATC classification mentioned above. In Table III, we can see how these drugs are classified by measuring their presence in percentages.

**Table III.** Analyzing Orphan Drugs by ATC Classification of Representative Drug Types.

| ATC classification | % orphan drugs |
| --- | --- |
| Antineoplastic agents | 68.57 |
| Dermatologicals | 5.71 |
| Nervous system | 5.71 |
| Antiparasitic products | 5.71 |
| Antiinfectives for systemic use | 5.71 |
| Cardiovascular system | 2.85 |
| Alimentary tract and metabolism | 1.42 |
| Systemic hormonal | 1.42 |
| Sensory organs | 1.42 |
| Musculo-skeletal system | 1.42 |

The results show that the most representative type of drugs as those used to treat neoplasms. This coincides with the most characteristic type of disease. Therefore, we want to make a final check to see if the repurposing of drugs in rare diseases occurs between diseases and drugs belonging to the same category. Fig 4 shows that the most abundant relationship between a rare disease and an orphan drug is around the neoplasm type. This result indicates that when there is a case of repurposing in rare diseases of the neoplasm type, this drug is also usually of this type. This pattern can be extrapolated to the rest of the categories since it seems that there is a direct relationship between the type of disease and drug being the same. We have also observed cases where this is not the case, for example, the relationship between neoplastic-type drugs and dermatological-type diseases that have an medium presence. But many cases follow the above-mentioned pattern.

**Fig 4.**
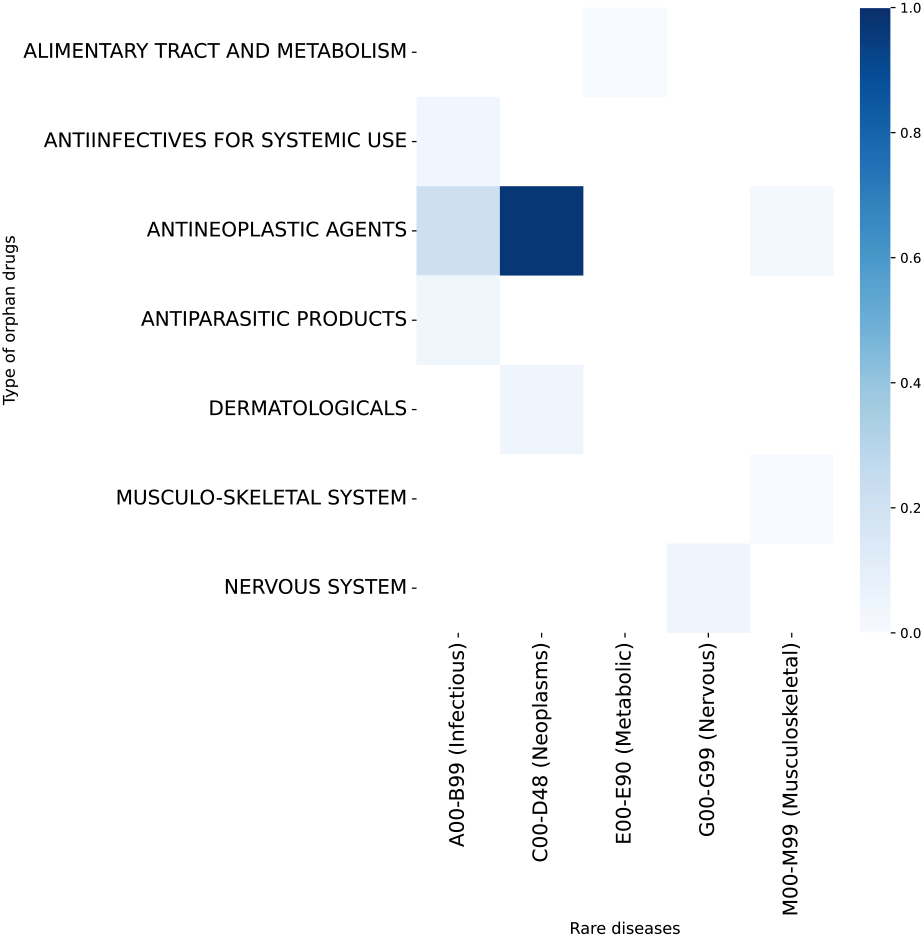
Relationship between the type of rare disease and the most common orphan drugs in the triples considered. The dark blue colour indicates a higher relationship between disease and drug, and the lighter colour indicates the opposite case.

Finally, the phenotypic similarity between the triplicate diseases was calculated. This similarity has been compared with the general data present in the DISNET platform. In Table IV, we can observe the results obtained when performing these comparisons and the statistical significance when applying the Mann-Whitney U test.

**Table IV.** PHENOTIPIC SIMILARITY BETWEEN RARE DISEASES AND DISEASES WAS CALCULATED USING THE JACCARD INDEX WITH THE RESULTING P-VALUE INDICATING STATISTICAL SIGNIFICANCE WHEN APPLYING THE MANN-WHITNEY U-TEST.

|  | *Phenotypic Similarity mean* | *P-value* |
| --- | --- | --- |
| Rare Disease – Disease | 0.206 | $2.73 \times 10^{-26}$ |
| DISNET | 0.054 |  |

If we look at the phenotypic similarity (Table V), we obtain a much higher value among the diseases that share orphan drugs by repurposing than if we compare it with the DISNET value. The differences between the two are statistically significant and the mean difference, in terms of percentages, is up to 10 times higher. This result indicates the importance of symptom sharing between diseases for which the same drug is used. This finding confirms what we have previously encountered in the scientific literature, where numerous instances of successful repurposing can be attributed to this phenomenon. Therefore, we can establish that the number of shared symptoms between two diseases, (in our case, rare diseases, since more than 80% of the triples are formed by two of these categories), is a very important starting point to consider a drug as a possible new treatment for another disease.

## V. Conclusions

Scientific research in the field of orphan drugs used to treat different rare diseases and repositioned drugs is very scarce. Despite the importance that has been gained in recent years in the different computational techniques of drug repurposing, there are very few studies that are focused on trying to find potential treatments for the multitude of rare diseases that are lacking. Employing the present study, it has been possible to reach different interesting conclusions that could contribute to alleviating this problem.

The main conclusion is that the repurposing of drugs in orphan drugs plays an important role that seems to become more relevant as the years go by. This will allow earlier and safer delivery of effective treatments for rare diseases that lack them. Focusing on the biological patterns, other conclusions of great interest were that most of the cases where a repurposing through an orphan drug occurs are between two rare diseases. Another of them was that the diseases shared by these drugs are usually of the same type. Regarding the type of drugs, it has been found that the most characteristic in the cases studied are neoplastic drugs used to treat neoplasms. In addition, it has been observed that in the cases of repurposing for rare diseases, the disease and the drug tend to be most of the same type. Focusing on phenotypic similarity, valuable results were obtained, where the similarity between rare and associated diseases was much higher than in general cases. This allows us to establish this similarity measure as a potential pattern for the search for new cases for repurposing orphan drugs.

## VI. Future steps and limitation

The main limitation of this study is the paucity of information regarding rare diseases. It is a poorly addressed field of research that lacks information in several aspects. Therefore, it has been difficult to obtain previous studies on the relationship between orphan drugs and drug repurposing. Another limitation was not being able to find CHEMBL and CUI for all the records obtained in the FDA database considered for this report.

As future lines of research, we would like to extend this study to the different world agencies that control the approval of drugs for rare diseases such as the European Medicines Agency (EMA) in the European Union. With a larger data source, we could establish clearer patterns that characterize rare diseases and allow us to advance further in this field of study. In addition, we would like to apply different NLP techniques on the names of chemical compounds that can help to identify the drugs they refer to associate their CHEMBL IDs to them. And, following along the same lines, apply these same NLP methods to the descriptions of the rare diseases found in the FDA database associated with these drugs to obtain their CUIS IDs. With these proposals, we intend to obtain a much more complete and representative study of all records related to orphan drugs, rare diseases, and drug repurposing worldwide.

## Acknowledgment

This research was funded by the projects “Data-driven drug repositioning applying graph neural networks (3DR-GNN)” (PID2021-122659OB-I00) from the Spanish Ministerio de Ciencia e Innovación, “Drug repurposing hypotheses through a data-driven approach (GRENADA)” (PDC2022-133173-I00) from the Spanish Ministerio de Ciencia e Innovación and MadridDataSpace4Pandemics, funded by Comunidad de Madrid (Consejería de Educación, Universidades, Ciencia y Portavocía) with FEDER funds as part of the response from the European Union to COVID-19 pandemia. Belen Otero Carrasco’s work is supported by “Formación de Personal Investigador” grant (FPI PRE2019-090912) as part of the project “DISNET (Creation and analysis of disease networks for drug repurposing from heterogeneous data sources applied to rare diseases)” (RTI2018-094576-A-I00) from the Spanish Ministerio de Ciencia, Innovación y Universidades. Andrea Álvarez Pérez was granted by Universidad Politécnica de Madrid and Banco Santander for a predoctoral ‘Programa Propio’ grant.

## Footnotes

1 https://disnet.ctb.upm.es/

2 https://www.accessdata.fda.gov/scripts/opdlisting/oopd/index.cfm

3 https://www.ebi.ac.uk/chembl/

4 https://www.nlm.nih.gov/research/umls/index.html

5 https://www.orpha.net/consor/cgi-bin/index.php

## References

[1] A. Angural et al., “Review: Understanding Rare Genetic Diseases in Low Resource Regions Like Jammu and Kashmir – India,” Front. Genet., vol. 11, 2020, Accessed: Mar. 10, 2023. [Online]. Available: https://www.frontiersin.org/articles/10.3389/fgene.2020.00415

[2] H. I. Roessler, N. V. A. M. Knoers, M. M. van Haelst, and G. van Haaften, “Drug Repurposing for Rare Diseases,” Trends Pharmacol. Sci., vol. 42, no. 4, pp. 255–267, Apr. 2021, doi:10.1016/j.tips.2021.01.003.

[3] S. Nguengang Wakap et al., “Estimating cumulative point prevalence of rare diseases: analysis of the Orphanet database,” Eur. J. Hum. Genet. EJHG, vol. 28, no. 2, pp. 165–173, Feb. 2020, doi:10.1038/s41431-019-0508-0.

[4] “RARE Disease Facts,” Global Genes. https://globalgenes.org/learn/rare-disease-facts/ (accessed Mar. 10, 2023).

[5] A. L. Crowe, A. J. McKnight, and H. McAneney, “Communication Needs for Individuals With Rare Diseases Within and Around the Healthcare System of Northern Ireland,” Front. Public Health, vol. 7, p. 236, Aug. 2019, doi:10.3389/fpubh.2019.00236.

[6] E. Kruger, P. McNiven, and D. Marsden, “Estimating the Prevalence of Rare Diseases: Long-Chain Fatty Acid Oxidation Disorders as an Illustrative Example,” Adv. Ther., vol. 39, no. 7, pp. 3361–3377, 2022, doi:10.1007/s12325-022-02186-2.

[7] X. Yan, S. He, and D. Dong, “Determining How Far an Adult Rare Disease Patient Needs to Travel for a Definitive Diagnosis: A Cross-Sectional Examination of the 2018 National Rare Disease Survey in China,” Int. J. Environ. Res. Public. Health, vol. 17, no. 5, Art. no. 5, Jan. 2020, doi:10.3390/ijerph17051757.

[8] S. Marwaha, J. W. Knowles, and E. A. Ashley, “A guide for the diagnosis of rare and undiagnosed disease: beyond the exome,” Genome Med., vol. 14, no. 1, p. 23, Feb. 2022, doi:10.1186/s13073-022-01026-w.

[9] H. A. [D-C.-24 Rep. Waxman, “Text - H.R.5238 - 97th Congress (1981-1982): Orphan Drug Act,” Apr. 01, 1983. http://www.congress.gov/ (accessed Mar. 10, 2023).

[10] R. Rodriguez-Monguio, T. Spargo, and E. Seoane-Vazquez, “Ethical imperatives of timely access to orphan drugs: is possible to reconcile economic incentives and patients’ health needs?,” Orphanet J. Rare Dis., vol. 12, no. 1, p. 1, Jan. 2017, doi:10.1186/s13023-016-0551-7.

[11] B. Delavan, R. Roberts, R. Huang, W. Bao, W. Tong, and Z. Liu, “Computational drug repositioning for rare diseases in the era of precision medicine,” Drug Discov. Today, vol. 23, no. 2, pp. 382– 394, Feb. 2018, doi:10.1016/j.drudis.2017.10.009.

[12] Regulation (EC) No 141/2000 of the European Parliament and of the Council of 16 December 1999 on orphan medicinal products, vol. 018. 1999. Accessed: Mar. 10, 2023. [Online]. Available: http://data.europa.eu/eli/reg/2000/141/oj/eng

[13] J. A. DiMasi, H. G. Grabowski, and R. W. Hansen, “Innovation in the pharmaceutical industry: New estimates of R&D costs,” J. Health Econ., vol. 47, pp. 20–33, May 2016, doi:10.1016/j.jhealeco.2016.01.012.

[14] S. Kraljevic, P. J. Stambrook, and K. Pavelic, “Accelerating drug discovery,” EMBO Rep., vol. 5, no. 9, pp. 837–842, Sep. 2004, doi:10.1038/sj.embor.7400236.

[15] B. Pereira Moreira et al., “Drug Repurposing and De Novo Drug Discovery of Protein Kinase Inhibitors as New Drugs against Schistosomiasis,” Molecules, vol. 27, no. 4, Art. no. 4, Jan. 2022, doi:10.3390/molecules27041414.

[16] D. Sardana, C. Zhu, M. Zhang, R. C. Gudivada, L. Yang, and A. G. Jegga, “Drug repositioning for orphan diseases,” Brief. Bioinform., vol. 12, no. 4, pp. 346–356, Jul. 2011, doi:10.1093/bib/bbr021.

[17] Y. Ko, “Computational Drug Repositioning: Current Progress and Challenges,” Appl. Sci., vol. 10, p. 5076, Jul. 2020, doi:10.3390/app10155076.

[18] N. Nosengo, “Can you teach old drugs new tricks?,” Nature, vol. 534, no. 7607, pp. 314–316, Jun. 2016, doi:10.1038/534314a.

[19] Y. Cha et al., “Drug repurposing from the perspective of pharmaceutical companies,” Br. J. Pharmacol., vol. 175, no. 2, pp. 168–180, Jan. 2018, doi:10.1111/bph.13798.

[20] Y. Cong and T. Endo, “Multi-Omics and Artificial Intelligence-Guided Drug Repositioning: Prospects, Challenges, and Lessons Learned from COVID-19,” OMICS J. Integr. Biol., vol. 26, no. 7, pp. 361–371, Jul. 2022, doi:10.1089/omi.2022.0068.

[21] R. Muthyala, “Orphan/rare drug discovery through drug repositioning,” Drug Discov. Today Ther. Strateg., vol. 8, no. 3, pp. 71–76, Dec. 2011, doi:10.1016/j.ddstr.2011.10.003.

[22] “Drug Therapy for Multiple Myeloma.” https://www.cancer.org/cancer/multiple-myeloma/treating/chemotherapy.html (accessed Mar. 10, 2023).

[23] F. Bellomo et al., “Drug Repurposing in Rare Diseases: An Integrative Study of Drug Screening and Transcriptomic Analysis in Nephropathic Cystinosis,” Int. J. Mol. Sci., vol. 22, no. 23, Art. no. 23, Jan. 2021, doi:10.3390/ijms222312829.

[24] R. G. Govindaraj, M. Naderi, M. Singha, J. Lemoine, and M. Brylinski, “Large-scale computational drug repositioning to find treatments for rare diseases,” NPJ Syst. Biol. Appl., vol. 4, p. 13, 2018, doi:10.1038/s41540-018-0050-7.

[25] M. Lotfi Shahreza, N. Ghadiri, and J. R. Green, “A computational drug repositioning method applied to rare diseases: Adrenocortical carcinoma,” Sci. Rep., vol. 10, no. 1, p. 8846, Jun. 2020, doi:10.1038/s41598-020-65658-x.

[26] G. Lagunes-García, A. Rodríguez-González, L. Prieto-Santamaría, E. P. García del Valle, M. Zanin, and E. Menasalvas-Ruiz, “DISNET: a framework for extracting phenotypic disease information from public sources,” PeerJ, vol. 8, Feb. 2020, doi:10.7717/peerj.8580.

[27] B. Otero-Carrasco, L. Prieto Santamaría, E. Ugarte Carro, J. P. Caraça-Valente Hernández, and A. Rodríguez-González, “Repositioning Drugs for Rare Diseases Based on Biological Features and Computational Approaches,” Healthc. Basel Switz., vol. 10, no. 9, p. 1784, Sep. 2022, doi:10.3390/healthcare10091784.

[28] M. Davies et al., “ChEMBL web services: streamlining access to drug discovery data and utilities,” Nucleic Acids Res., vol. 43, no. Web Server issue, pp. W612–W620, Jul. 2015, doi:10.1093/nar/gkv352.

[29] A. S. Brown and C. J. Patel, “A standard database for drug repositioning,” Sci. Data, vol. 4, no. 1, Art. no. 1, Mar. 2017, doi:10.1038/sdata.2017.29.

[30] “International Classification of Diseases (ICD).” https://www.who.int/standards/classifications/classification-of-diseases (accessed Nov. 03, 2022).

[31] “Anatomical Therapeutic Chemical (ATC) Classification.” https://www.who.int/tools/atc-ddd-toolkit/atc-classification (accessed Nov. 18, 2022).

